# A lethal ORC ATPase mutation is suppressed by alterations in ORC and RNA Pol II transcription components

**DOI:** 10.64898/2026.05.01.722367

**Authors:** Luis E. Martinez-Rodriguez, Stephen P. Bell

**Author notes:** Corresponding author: (SPB).

## Abstract

The origin recognition complex (ORC) selects origins of replication and directs the loading of the Mcm2-7 replicative helicase at these sites. Five of the six ORC subunits are related to the AAA+ family of ATPases. Although functions for ATP hydrolysis by Cdc6 and the Mcm2-7 complex have been described, the essential role of ORC ATP hydrolysis remains unclear. We performed a genetic screen in *Saccharomyces cerevisiae* for suppressors of the lethal phenotype of the *orc4-R267A* allele, which disrupts ORC ATP hydrolysis *in vitro*. We identified six causative mutations, five of which are distributed across different ORC subunits. The suppressor mutations in Orc1 and Orc4, but not the other ORC subunits, increase the *in vitro* helicase loading activity of ATPase-defective ORC (ORC4R). Allele specificity studies showed the alleles specifically suppress defects at ATPase interfaces within the ORC-Cdc6 complex. The sixth allele is a mutation in *TOA2*, a subunit of the TFIIA general transcription factor. Mutations in the general transcription factors TBP and TFIIB, and the large subunit of RNA Polymerase II also suppressed the *orc4-R267A* lethality, suggesting that reducing transcription is sufficient for suppression. Our study identifies multiple ways to suppress the lethal phenotype of an ATPase defective ORC allele and reveals a connection between ORC ATP hydrolysis and transcription.

## Introduction

Eukaryotic DNA replication initiation is a tightly regulated process that assembles multi-enzyme machines at origins of replication. This process is separated into two cell-cycle-regulated events. During the G1 phase of the cell cycle, two copies of the hetero-hexameric replicative DNA helicase, the Mcm2-7 complex, are loaded onto each origin as an inactive head-to-head “double hexamer”. Importantly, this event, referred to as helicase loading or origin licensing, marks all potential origins of replication. When cells enter S phase, loaded helicases are activated while *de novo* helicase loading is inhibited for the rest of the cell cycle [1]. This temporal regulation of helicase loading and activation prevents repeated origin licensing and ensures that the genome is replicated once per cell division [2].

Helicase loading is a multistep process requiring several proteins: the origin recognition complex (ORC), Cdc6, Cdt1, and the Mcm2-7 complex. ORC, a six-subunit complex, initiates helicase loading by recognizing a conserved sequence (the ARS consensus sequence or ACS) found at each budding yeast origin of replication in an ATP-dependent manner [3]. Cdc6 binds DNA-bound ORC between Orc1 and Orc2, to complete a six-protein ring that encircles DNA [4,5]. This assembly recruits an Mcm2-7 hexamer in complex with Cdt1, forming a short-lived intermediate called the ORC-Cdc6-Cdt1-Mcm2-7 (OCCM) complex [6,7]. Subsequently, Cdc6 then Cdt1 release from the DNA while ORC repositions to the opposite face of the first Mcm2-7 hexamer [8–11]. The later steps of this process are repeated in a quasi-symmetric manner to recruit and load the second Mcm2-7 complex, resulting in a Mcm2-7 double hexamer that encircles DNA.

Twelve of the fourteen proteins involved in helicase loading are related to the ATPases-associated-with-various-activities (AAA+) family of proteins: Orc1-5 subunits, Cdc6, and all six Mcm subunits. AAA+ ATPase proteins use the energy of ATP binding and hydrolysis to remodel or translocate on macromolecules [12]. AAA+ domains form multimers with the ATPase active site at the interface between two subunits. One subunit contributes the Walker-A and -B motifs, which are key components of the ATP binding site. The adjacent subunit completes the ATPase by providing an arginine-finger motif that stimulates ATP hydrolysis. Mutations in these motifs frequently distinguish the role of ATP binding (Walker-A mutations) from ATP hydrolysis (arginine-finger or Walker-B mutations) [13].

The functions of ATP hydrolysis by Cdc6 and MCM have been characterized. Each of the AAA+ motifs in Cdc6 and all but one of the Mcm2-7 ATPase interfaces are essential for viability [14–17]. Cdc6 ATP hydrolysis is not required for ORC-DNA binding or helicase loading *in vitro* or *in vivo*. Instead, Cdc6 ATP hydrolysis releases both Cdc6 from the OCCM intermediate and defective Mcm2-7 complexes [16,18,19]. Mcm2-7 ATP hydrolysis drives many of the events in DNA replication initiation. All Mcm2-7 ATPase mutants show defects in helicase loading [10,16,17,20]. After helicase loading, MCM ATP hydrolysis also promotes efficient recruitment of helicase activators, initiates origin melting, and powers DNA template unwinding during DNA replication [16,21–24].

In contrast to Cdc6 and Mcm2-7, the role of ORC ATP hydrolysis is unclear. The only ORC ATPase site is located at the Orc1-Orc4 interface and mutation of the AAA+ motifs at this interface are lethal *in vivo* [25,26]. *In vitro*, mutation of the Walker-A motif abolishes ATP binding and inhibits sequence-specific DNA binding [26]. In contrast, mutation of the arginine finger in Orc4 (Orc4 R267A, also referred to as ORC4R) abolishes ATP hydrolysis but does not prevent DNA binding [25]. Overexpression of ORC4R protein prevents Mcm2-7 loading *in vivo* but the ORC ATPase-deficient protein promotes helicase loading *in vitro* [17,25]. ORC ATP hydrolysis has also been implicated in the establishment of chromatin organization at origins but the mechanism of this function remains unclear [27].

Inactivation of Orc1 or Orc2 *in vivo* is sufficient to prevent Mcm2-7 origin association [28] and cell cycle execution point analysis shows that ORC function is required exclusively during G1 [29]. Moreover, mutations in the Mcm2-7 complex that specifically disrupt Mcm2-7 recruitment to ORC-Cdc6-DNA complex abolish loading and cause lethality [18]. These studies strongly suggest that ORC’s essential function is recruiting and loading the Mcm2-7 complex. Because the ORC4R protein can promote helicase loading *in vitro* but the *orc4-R267A* allele is lethal *in vivo*, how ORC ATP hydrolysis contributes to helicase loading remains unclear.

Transcription impacts DNA replication in several ways beyond its role in the synthesis of the DNA-replication proteins [30]. First, transcription through an origin has been shown to inactive origins by displacing the Mcm2-7 helicase [31,32]. Second, because RNA polymerase progression requires disassembly and reassembly of nucleosomes, it shapes the chromatin environment throughout the genome, including origins [33,34]. Finally, transcribing RNA polymerase can physically collide with the replisome during S phase, which results in collapse of the one of the two synthesis machines [35–38].

Here, we use a combination of genetic and biochemical experiments to address the role of ORC ATP hydrolysis during DNA replication initiation. Using a genetic screen for suppressors of the lethal phenotype of the *orc4-R267A* ORC ATPase mutant, we identified six suppressing mutations. Five of the mutations are in ORC subunits and one suppressor is in *TOA2*, a subunit of the general transcription factor TFIIA. The *orc1* and *orc4* suppressor alleles increase helicase-loading activity *in vitro*. The remaining ORC mutations did not enhance the biochemical activities of ORC4R. Allele specificity studies showed these alleles suppressed a variety of mutations in the ORC-Cdc6 AAA+, suggesting a connection between the two ATPase sites. Cdc6 overexpression was sufficient to suppress the lethal phenotype of the *orc4-R267A* allele. Finally, alleles in subunits of general transcription factors, such as TFIIB and TBP, and RNA polymerase II also showed partial suppression of the *orc4-R267A* allele. Our study identified multiple ways to suppress the lethal phenotype of an ATPase defective ORC mutant, establishing potential connections between the two ORC-Cdc6 ATPase sites and RNA transcription.

## Results

### A screen for suppressors of a lethal ORC ATPase mutant

Because ORC containing the Orc4-R267A ATPase mutation (ORC4R) is not defective for helicase loading *in vitro* [17], we took a genetic approach to investigate the essential function of ORC ATP hydrolysis. To this end, we performed a screen for suppressors of the lethal *orc4-R267A* phenotype (Fig 1A). We mutagenized a strain with *orc4-R267A* integrated into the genome that was kept alive by a counter-selectable, complementing plasmid that included *ORC4*, *URA3,* and *ADE2* genes. We identified strains with suppressor mutations by their ability to grow in the absence of the plasmid as determined by growth on 5-fluorouracil (5-FOA, indicating loss of *URA3*) and red colony color (indicating loss of *ADE2*). The use of two markers for plasmid loss eliminated false positives due to a *URA3* or *ADE2* mutation. We screened ∼10^8^ cells and isolated 72 red, 5-FOA-resistant colonies. We focused on 24 strains that grew at a similar rate to the starting strain in YPD media. Using tetrad analysis of strains backcrossed to the starting strain, we identified eight strains whose suppressor phenotype showed 2:2 segregation, consistent with a single allele being responsible for the phenotype.

**Fig 1.**
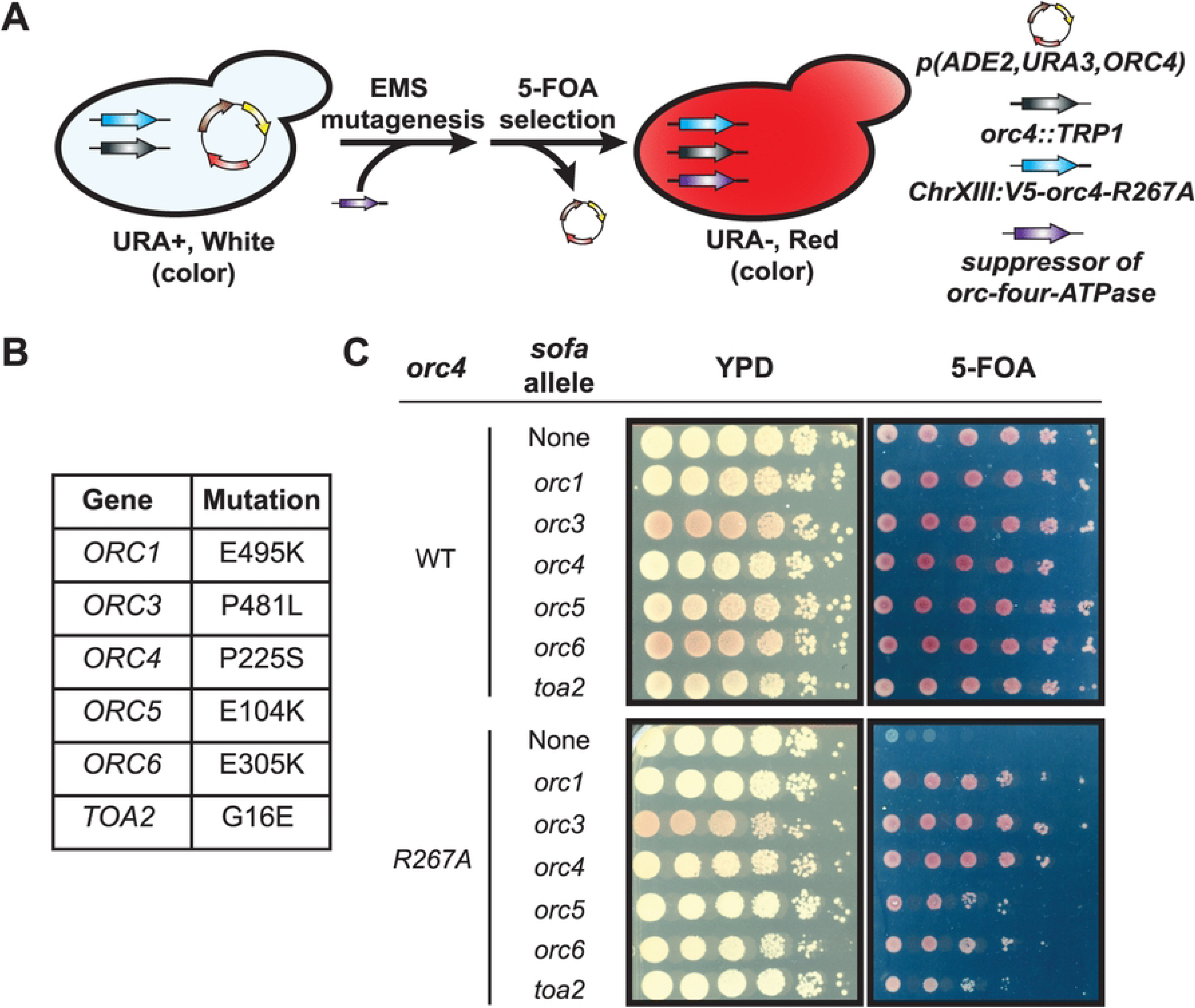
Screen for suppressors of the *orc4-R267A* lethal phenotype. A) An *orc4-R267A* strain that was kept alive by a plasmid containing *ORC4* was treated with ethyl methane sulfonate (EMS) to isolate suppressors of the lethal phenotype of the *orc4-R267A* allele. Only strains that acquire suppressors can lose the plasmid. The desired strains turn red and grow on 5-Fluorouracil (5-FOA) because they lose the *ADE2* and *URA3* genes on the plasmid. B) Table of *orc4-R267A* suppressor mutations identified in the screen described in A. C) Ten-fold serial dilutions of strains that have *ORC4* on a counter-selectable plasmid and either *ORC4* or *orc4-R267A* allele integrated into the genome. The presence of <u>s</u>uppressors-of-<u>o</u>rc-four-<u>A</u>TPase alleles are indicated. Some strains are red prior to 5-FOA selection because the suppressor allele allowed plasmid loss during strain construction.

To identify the causative mutation for each suppressor phenotype, we used pooled linkage analysis with whole-genome sequencing [39,40]. Six of the eight strains identified a single missense mutation likely to be responsible for the observed suppression (Fig 1B). We identified mutations in five of the six ORC subunits (all but *ORC2*). Interestingly, we also identified a mutation in *TOA2*, a subunit of the RNA Pol II general transcription factor TFIIA [41]. We will refer to the suppressor alleles by their gene name followed by <u>s</u>uppressors-of-<u>o</u>rc-four-<u>A</u>TPase (*sofa*). For example, the *orc4* suppressor allele will be referred to as *orc4-sofa*.

To confirm that the identified mutations are sufficient to suppress the lethal phenotype of *orc4-R267A*, we used CRISPR-Cas9 to regenerate each mutation in the starting-strain background [42,43]. For each potential suppressor, we asked if the resulting strain could grow if the only *ORC4* allele present was *orc4-R267A* (on 5-FOA). All the mutations restored growth on 5-FOA, although the extent of suppression differed (Fig 1C). The *orc1-, orc3-,* and *orc4-sofa* alleles restored growth to near WT levels, whereas the *orc5-, orc6-,* and *toa2-sofa* alleles resulted in slower growth. None of the suppressor mutations had a growth defect when regenerated in an *ORC4* background (Fig 1C). Furthermore, in the *orc4-R267A* background but not the WT background, all the six *sofa* alleles were sensitive to high temperature (37 °C) while four of the six, the *orc1*, *orc5*, *orc6* and *toa2* alleles, were sensitive to 200 mM hydroxyurea (S1A and S1B Figs).

### Location of the suppressor mutations in 3D structures

As a first step to investigate the mechanism of suppression for the identified mutations, we located the site of the ORC *sofa* mutations in the 3D-structure of an ORC-Cdc6-DNA complex and the location of the Toa2 mutation in the TFIIA-TBP-DNA complex (Fig 2) [4,44,45]. None of the ORC mutations are in amino acids that contact the ATP molecule at the Orc1-Orc4 ATP hydrolysis site. Although not adjacent to the ATP, the orc1-*sofa* mutation (E495K) is on the same helix as the Walker-A lysine, raising the possibility that it could indirectly impact the ATP hydrolysis site. Two of the mutations in ORC are part of regions or domains that could impact DNA binding indirectly. The *orc5-sofa* mutation (E104K) is 9 Å from the DNA, near a positively charged Orc1 region that interacts with the minor groove of the DNA. The mutated residue in *orc6-sofa* (E305K) is in the Orc6 C-terminal domain that interacts with the bent region of the ORC-bound DNA. Finally, the *orc3-sofa* mutation (P481L) is located at an interface between Orc2, Orc3, Orc5, and Orc6 while the *orc4-sofa* mutation (P225S) is in a linker region of the AAA+ Rossmann fold. Although the location of the mutated residues on the ORC-Cdc6 structure doesn’t provide a clear mechanism for suppression, the diverse locations of these residues raise the possibility that they impact multiple aspects of ORC function. We also inspected the location of the mutated amino acids in two other structures of complexes relevant to helicase loading, the OCCM and MO complexes [6,11], but found no changes in the location of the mutated amino acids that might explain the suppression.

**Fig 2.**
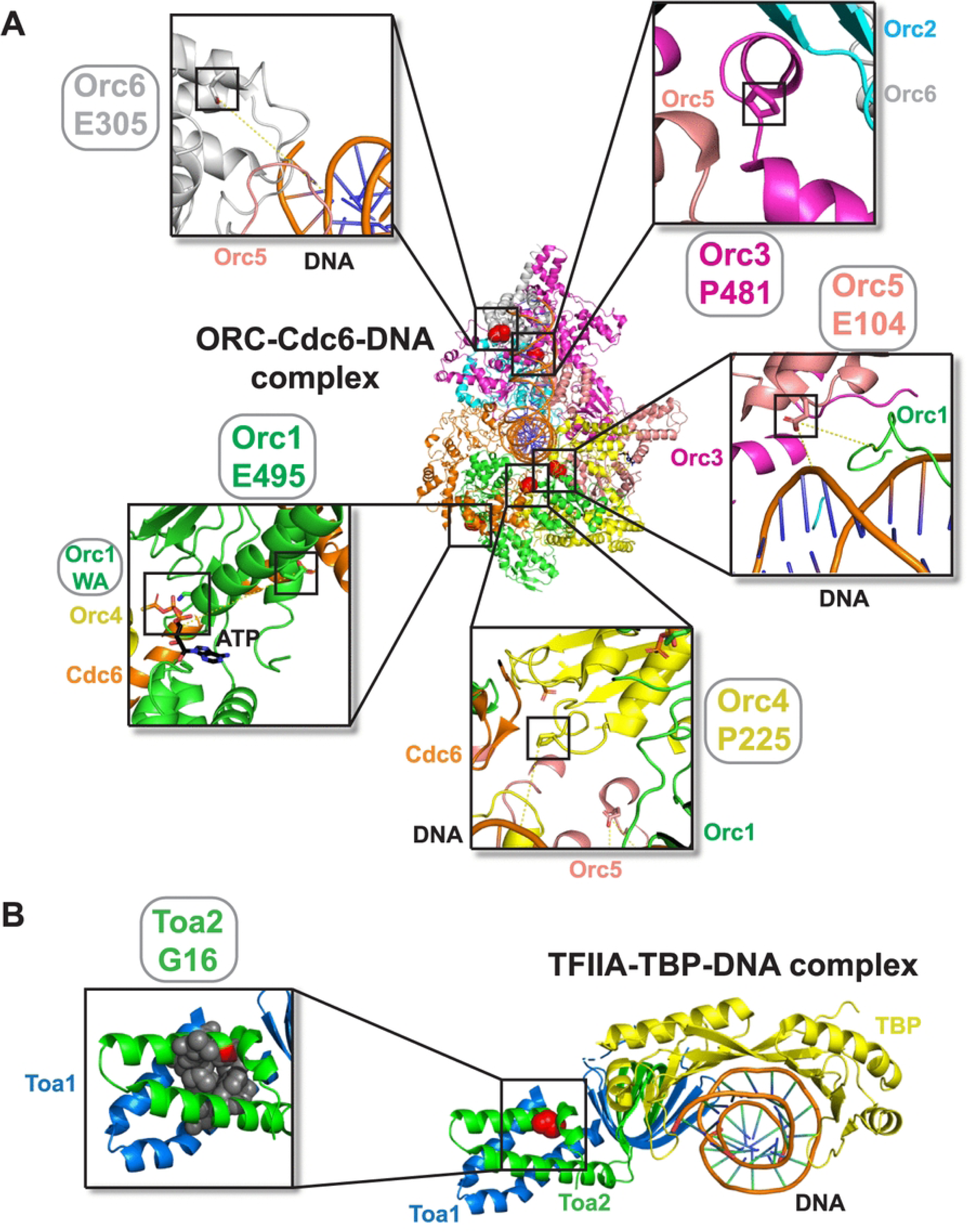
Location of residues mutated in orc4-R267A-suppressor alleles in 3D protein structures. A) The ORC-Cdc6 complex bound to DNA (PDB: 7MCA). For each allele, the location is marked in the full structure by a black box, and an enlarged version of the same region is shown. The mutated residue and respective subunits are indicated. ATP, DNA, other protein subunits or AAA+ motif residues visible in the enlarged region are also labeled. B) The TFIIA-TBP-DNA complex bound to DNA (PDB: 1NH2). In the structure, the mutated Toa2 residue (G16) is shown as red spheres. An enlarged version of the region including the mutated residue is shown. Hydrophobic residues on both Toa1 and Toa2 near the mutated residue are shown as grey spheres.

We performed multiple sequence alignment of the ORC protein sequences that were mutated in the suppressors [46–48] (S2 and S3 Figs). All the residues are conserved within *Saccharomycetaceae* except for the Orc6 residue. None of the residues are conserved between yeast and other metazoan model organisms, although in some cases there is evidence for chemical conservation. For example, the residue mutated in Orc1, Orc5, and Orc6, appears to be consistently polar across all species (S2A, S3A and S3B Figs). Similarly, although the proline residue mutated in Orc3 is not conserved, in most organisms there is another proline close by that is conserved and could serve the same function (S2B Fig). Because the mutated residues are primarily conserved within *Saccharomycetaceae*, the residues are likely to serve a specific function within these organisms.

The Toa2 mutation was in a glycine at the end of the first alpha helix of the Toa2 protein. According to a Ramachandran plot, the glycine conformation was not in a glycine-exclusive region [49,50]. Instead, the residue was close to a hydrophobic core formed between Toa1 and Toa2. Unlike the ORC mutated residues, the mutated *TOA2* residue is conserved across all organisms examined (S3C Fig). Based on the location, replacement of glycine with a glutamate likely destabilizes hydrophobic interactions between Toa2-Toa1 in the TFIIA complex.

### The *sofa* mutations have different effects on the biochemical activities of ORC4R protein

The simplest mechanism of suppression is that the *sofa* mutations restore ORC ATPase activity. We purified ORC with the Orc4 arginine-finger mutation (ORC4R) [25] as well as ORC4R including each one of the five *sofa* mutations in the ORC subunits (e.g., ORC4R^Orc1-*sofa*^). We tested each of these complexes for the ability to hydrolyze ATP relative to WT ORC (Fig 3A). None of the mutations restored ORC ATPase activity, ruling out this effect as a mechanism of suppression.

**Fig 3.**
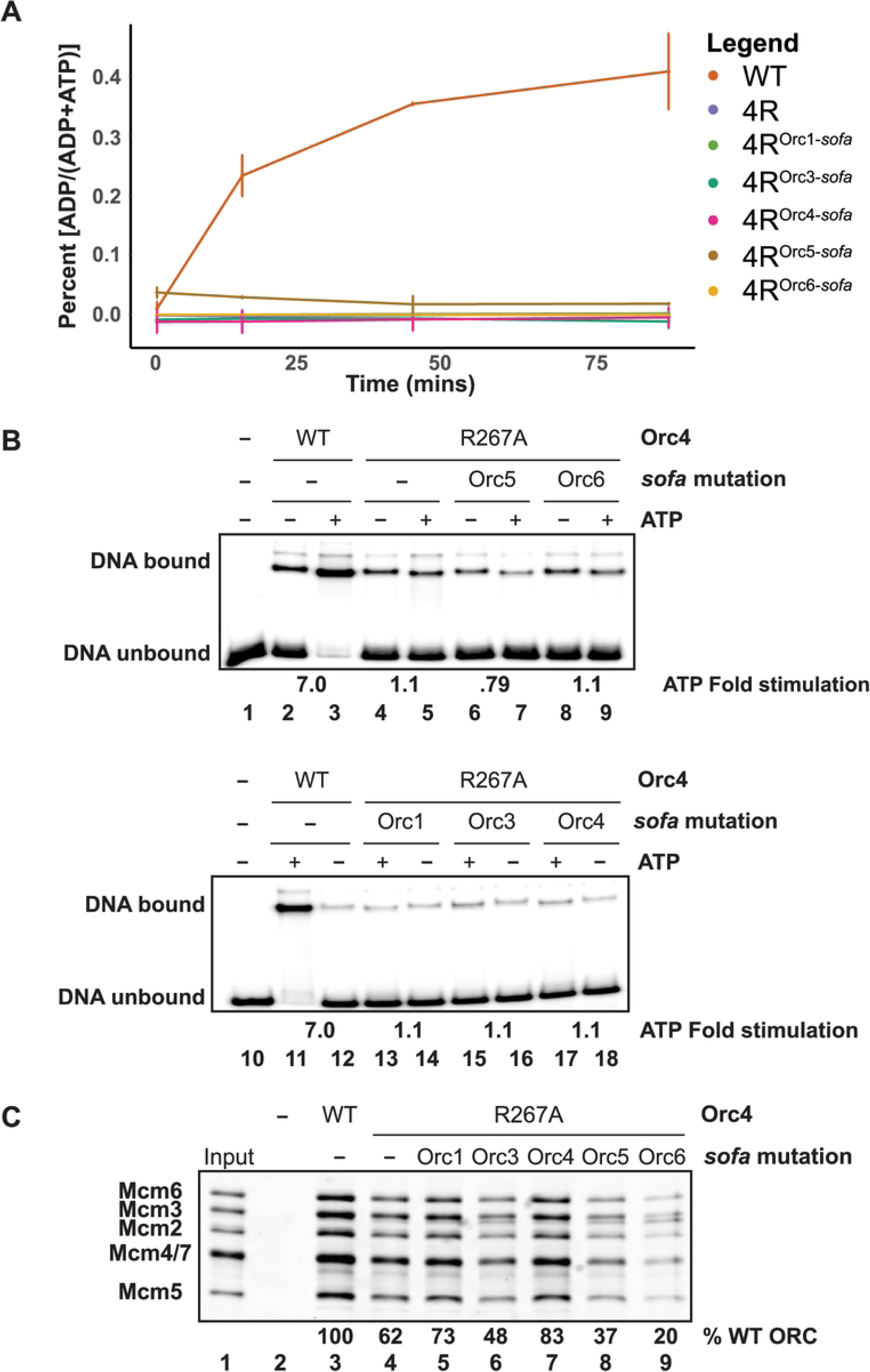
Biochemical effects of suppressor mutations on the biochemical activities of the ORC4R protein. A) ATPase activity of WT ORC and ORC4R protein with or without an additional suppressor mutation. Hydrolysis was monitored using radioactive ATP. Percent hydrolysis (ADP / (ADP+ATP), arbitrary units) was determined for WT, ORC4R, and ORC4R^sofa^ proteins over a time course. B) Electrophoretic-mobility shift assay to determine *ARS1* DNA binding of mutant protein complexes. Fold stimulation was calculated by dividing the DNA bound signal in the presence of ATP by the DNA bound signal in the absence of ATP. C) *In vitro* helicase-loading assay for WT and ORC4R protein without or with the indicated suppressor mutation.

Two of the residues mutated in the suppressor alleles were in regions near the DNA molecule in the ORC-Cdc6-DNA structure raising the possibility that they impacted DNA binding. We used an electrophoretic-mobility shift assay (EMSA) [51] to assess the impact of the *sofa* mutations on ORC4R DNA-binding (Fig 3B) with or without ATP. In contrast to previous footprinting-based assays [25], we observed clear DNA-binding defects for ORC4R. Unlike the seven-fold stimulation seen in the wild-type ORC protein, ORC4R showed no increase in binding upon ATP addition. Importantly, none of the ORC subunit *sofa* mutations tested led to an increase in ORC4R DNA binding with or without ATP. Thus, the suppressor mutations do not restore the *in vitro* DNA-binding defect of ORC4R.

Finally, we tested the impact of the *sofa* mutations on ORC’s essential function using a reconstituted helicase-loading assay [52,53] (Fig 3C). The suppressors had different effects on ORC4R loading activity. The ORC4R^Orc1-*sofa*^ and ORC4R^Orc4-*sofa*^ proteins showed a 16 and 28 % increase, respectively, in loading activity relative to ORC4R but neither reached WT levels (Fig 3C). In contrast, the ORC4R^Orc3-*sofa*^, ORC4R^Orc5-*sofa*^, and the ORC4R^Orc6-*sofa*^ proteins each decreased helicase-loading activity (Fig 3C). Under the higher-salt conditions that further reduced ORC4R loading relative to WT ORC, the ORC4R^Orc1-*sofa*^ and ORC4R^Orc4-*sofa*^ proteins showed 9- and 10-fold higher loading (S4A Fig). Even though the loading of the ORC4R protein with the Orc4-*sofa* mutation did not reach WT levels, the protein’s loading activity was consistently ∼34 to∼78% higher than the ORC4R protein across the salt titration (S4B Fig). These findings suggest that the *orc-sofa* mutations are likely to restore ORC function by different mechanisms. To further characterize the ORC4R defect (Fig 3B and 3C), we investigated its DNA-binding and helicase-loading activities in more detail.

The ORC4R protein had low amounts of DNA-binding, regardless of nucleotide presence or the addition of the *sofa* mutations (Fig 3B). This finding contrasted with previous findings suggesting that ATP hydrolysis is not required for ORC-DNA binding [3,26] and that the Orc4-R267A mutation did not cause defects in a footprinting-based DNA-binding assay [25]. To confirm that ATP hydrolysis was not required for DNA binding, we used the EMSA assay to test if ORC could bind DNA in the presence of ATPγS, a slowly hydrolyzable ATP analog (S5A Fig). ORC DNA binding was similar for WT ORC in the presence of either ATP or ATPγS, confirming that ATP hydrolysis is not required for DNA binding in this assay. We included the Walker-A defective ORC protein complex, which contains a mutation at lysine 485 of the Orc1 subunit (ORC1K), as a control and observed a similar DNA-binding defect to the ORC4R protein. Although not ATP-dependent, the ORC4R DNA-binding activity is sequence-specific (S5B Fig). Since we see small amounts of DNA binding for all the ORC complexes in the absence of ATP, this suggests that ATP binding stimulates ORC DNA binding as opposed to being strictly necessary.

Even though the ORC4R protein showed a DNA binding defect, it retained ∼60% of the wild-type protein’s helicase-loading activity (Fig 3C), consistent with previous studies [17]. To ensure we did not overcompensate for the weak DNA-binding of the ORC4R protein by adding high concentrations of ORC, we titrated the WT, ORC4R, and ORC1K proteins (S5C Fig). At lower concentrations, ORC4R retained ∼34% activity relative to the WT protein whereas ORC1K retained ∼15% activity. To determine whether the residual ORC4R loading activity reflected a different DNA binding mode, we titrated the salt concentrations in the loading reaction (S4B Fig). Previous studies showed that higher salt concentrations reveal the sequence-specific nature of helicase loading, as low-salt conditions allow ACS-independent helicase loading [54]. The ORC4R protein was sensitive to increased salt concentrations, consistent with a DNA-binding defect (S5D, S4A and S4B Figs), and loading remained origin sequence-specific at lower salt concentration (250 mM), regardless of the *orc4-sofa* mutation (S5D Fig). Although ORC4R and ORC1K show similar DNA-binding defects, ORC4R retains two- to three-fold more helicase-loading activity (S5C Fig), suggesting that the Orc4 arginine-finger mutation preserves some ATP-dependent activities required for loading that are lost in the ATP-binding-deficient ORC1K. Given that several of the *sofa* mutations further reduce ORC4R’s loading activity yet still suppress lethality *in vivo* (Fig 1 and 3, S4A Fig), these findings suggest that suppression is not solely mediated by restoring ORC biochemical function.

### Allele specificity of the *sofa* mutations

We next investigated the allele-specificity of the *sofa* mutations. First, we asked if the *sofa* mutations also suppressed a mutation that eliminated ATP binding (as opposed to ATP hydrolysis) in the same ATPase active site. ATP binding at this site is important for ORC origin-DNA binding *in vitro* and mutations that impair ATP binding at this site are lethal *in vivo* [26] (Fig 4). Using a similar approach described in Figure 1A, we asked if each *sofa* mutation could suppress an ATP-binding-defective *ORC1* mutation, *orc1-K485T* (Fig 4A). Interestingly, all the mutations except the *orc1-sofa* suppressed the lethal phenotype of the *orc1-K485T* allele.

**Fig 4.**
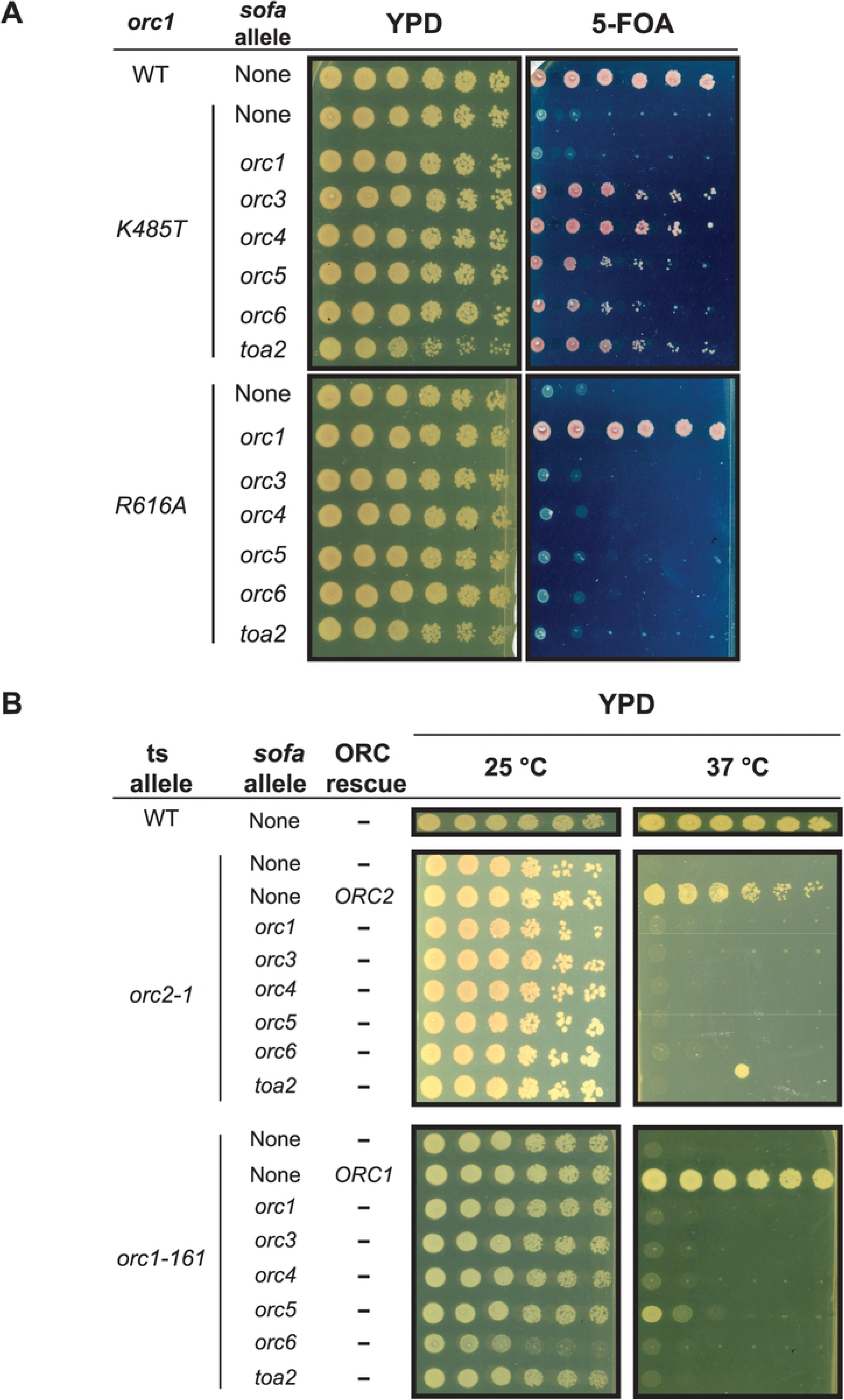
Allele specificity of the *sofa* alleles. A) Ten-fold serial dilutions of *ORC1, orc1-K485T*, or *orc1-R616A* plasmid-shuffle strains without and with each of the six *sofa* alleles. Strains were incubated 2-3 days at 30°C. Lysine 485 is in the Orc1 Walker-A motif at the Orc1/Orc4 ATPase interface. Arginine 616 is in the Arginine-finger motif of Orc1 at the Cdc6/Orc1 ATPase interface. B) Ten-fold serial dilutions of *orc2-1* and *orc1-161* strains without and with additional *sofa* alleles. Strains were incubated 2-3 days at 25°C and 37°C.

We also tested the ability of the *sofa* mutations to suppress a mutation in a second essential ATPase formed between the Orc1 and Cdc6 proteins [4,19]. The Cdc6-ORC-DNA complex is responsible for recruiting the Mcm2-7/Cdt1 complex and a second ATPase site forms when Cdc6 binds the ORC-DNA complex. Consistent with the importance of this ATPase, a mutation in Orc1 that eliminates the arginine finger that is essential for ATP hydrolysis at the Cdc6/Orc1 interface (*orc1-R616A*) is lethal [4] (Fig 4A). Interestingly, this allele showed a complete reversal of the suppression pattern for *orc1-K485T*, with only *orc1-sofa* suppressing the lethal phenotype of this Cdc6-Orc1 ATPase mutant. The ability of *orc1-sofa* to suppress ATPase mutations in the Orc1-Orc4 and Cdc6-Orc1 interfaces suggests a functional connection between these ATPases.

As a final test of allele specificity of the *sofa* alleles, we determined if the *sofa* mutations could suppress either of two temperature-sensitive (ts) ORC mutants: *orc1-161* and *orc2-1* [28,29,55] (Fig 4B). Both alleles affect the stability of the ORC complex *in vivo* [29,56]. Only the *orc5-sofa* weakly suppressed *orc1-161*. Thus, the *sofa* mutations are largely specific to mutations that affect the Orc1-Orc4 ATPase interface.

### *CDC6* overexpression suppresses the lethal phenotypes of orc4-R267A *and* orc1-K485T

Cdc6 binding between Orc1 and Orc2 stabilizes ORC on DNA [10,57,57,58], providing a possible mechanism to suppress the ORC4R DNA binding defect. To test if Cdc6 overexpression suppresses the lethal phenotype of the AAA+ motif mutant alleles, we overexpressed Cdc6 using a GAL-inducible promoter in *orc1-K485T*, *orc1-R616A*, and *orc4-R267A* plasmid-shuffle strains (Fig 5). GAL overexpression of Cdc6 suppressed the lethal phenotype of both the *orc4-R267A* and the *orc1-K485T* alleles but not the *orc1-R616A* allele, suggesting that Cdc6 overexpression can compensate for a defect in the Orc1-Orc4 ATPase but not defects in the Cdc6-Orc1 ATPase. This suppression also provides further support for a functional connection between these two ATPases.

**Fig 5.**
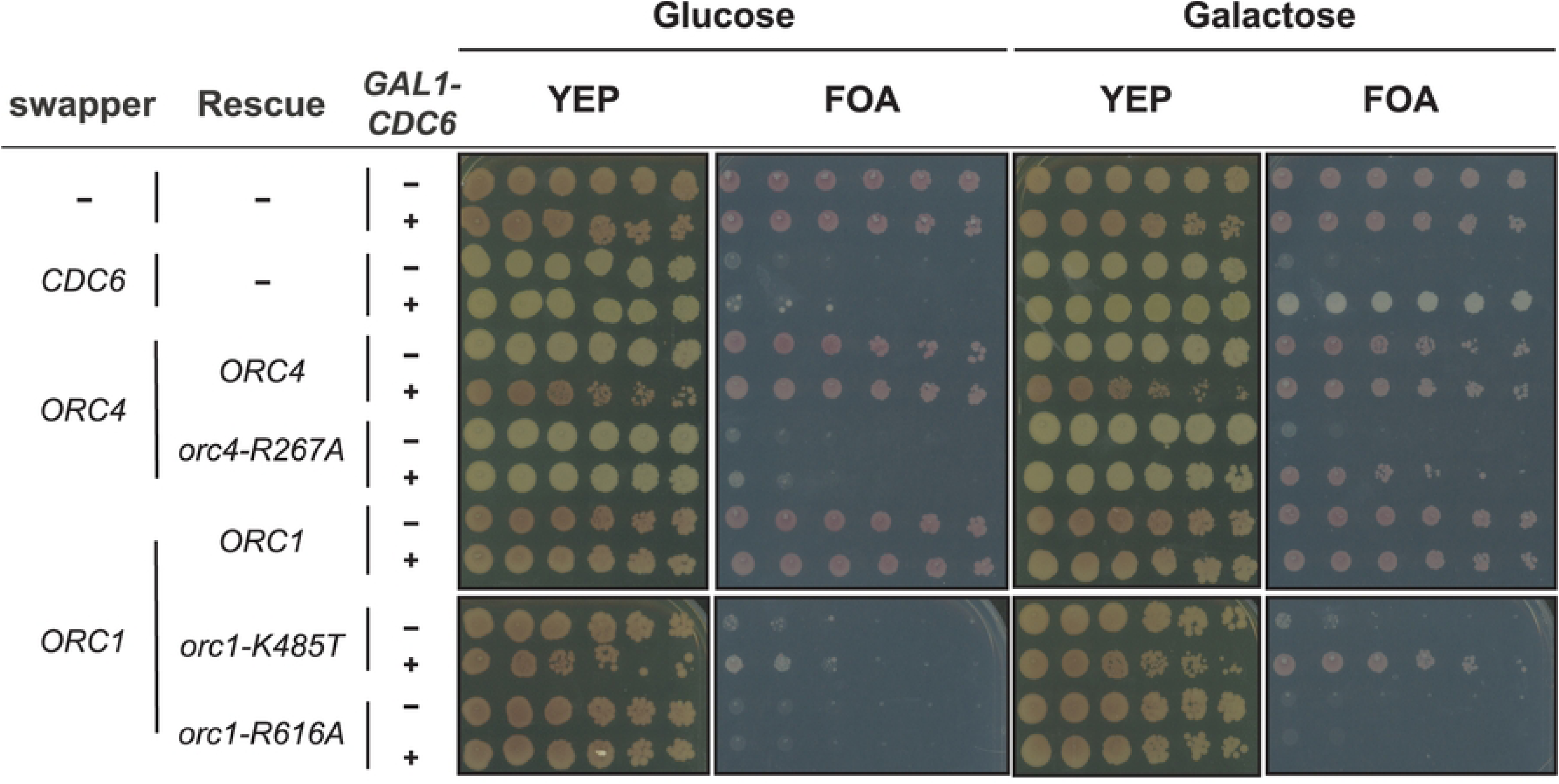
Galactose-induced CDC6 overexpression of *ORC4-* and *ORC1-* plasmid-shuffle strains. Ten-fold serial dilutions of *ORC4*, *orc4-R267A*, *ORC1*, *orc1-K485T*, and *orc1-R616A* plasmid-shuffle strains with or without induction of the *GAL-CDC6* allele. A control strain with a *CDC6* deletion covered by plasmid with wild-type Cdc6 is also shown (rows 3 and 4). All strains were grown overnight in galactose, plated, and incubated at 30°C for 3 days.

### Reduced RNA Pol II transcription suppresses the lethal phenotype of *orc4-R267A*

Toa2 interacts with Toa1 to form the general transcription factor TFIIA that binds TATA-binding protein (TBP) to stabilize its interaction with DNA [41]. Based on the location of the Toa2 mutation in the TFIIA-TBP-DNA structure, the *toa2-sofa* mutation likely disrupts the TFIIA complex (Fig 2B). We hypothesized that other mutations that destabilize the TFIIA complex or reduce its interaction with TBP should serve as suppressor mutations. To test this hypothesis, we tested an allele that disrupts TFIIA-TBP interaction *in vitro, toa1-V251F* [44,59,60] (Fig 6). Consistent with our hypothesis, the *toa1-V251F* allele also suppresses the lethal phenotype of the *orc4-R267A* allele. The suppressor phenotypes of the toa1 and toa2 mutations, together with their locations in the structure, suggest that disruption of the TFIIA complex underlies the suppression.

**Fig 6.**
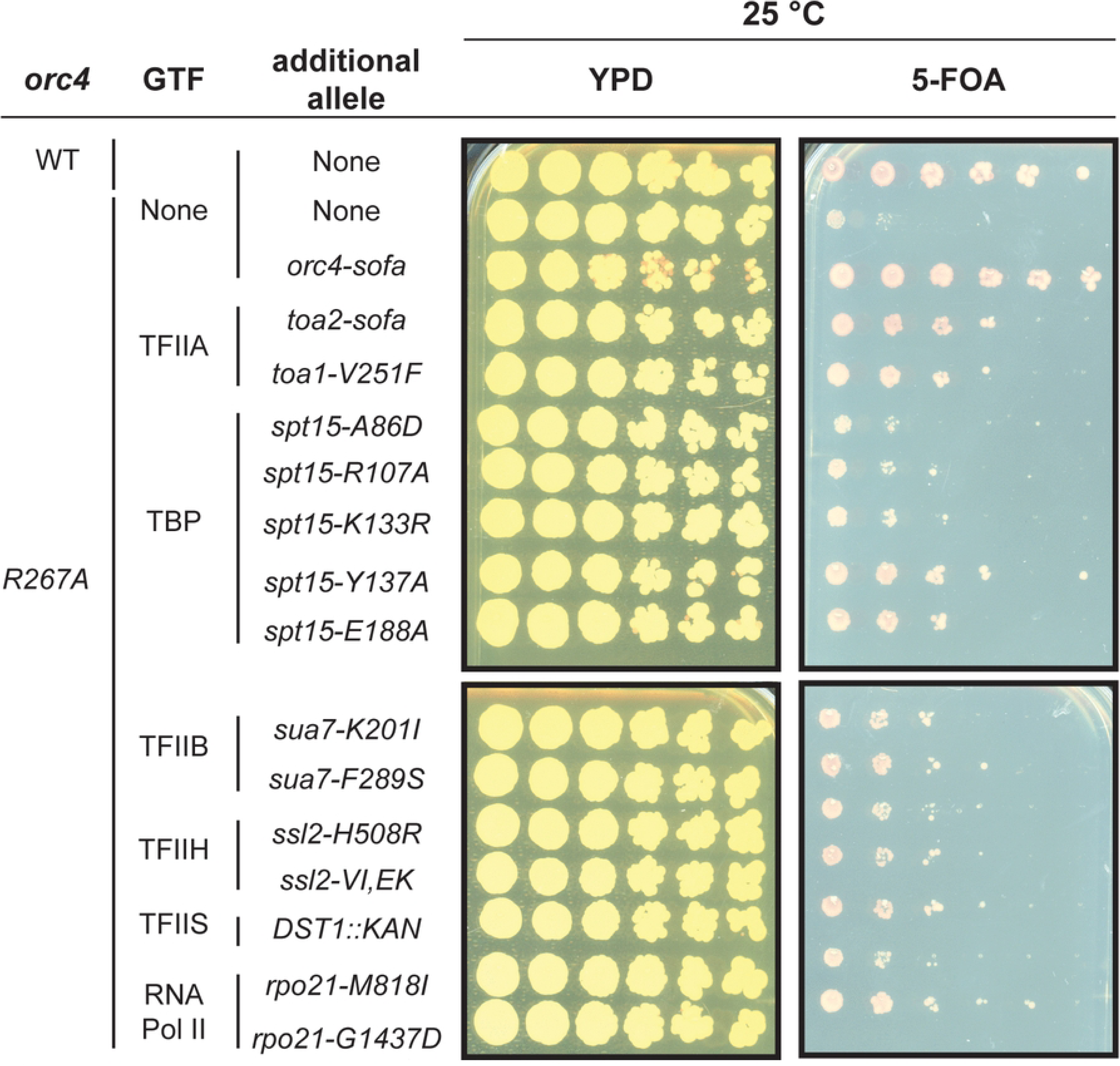
Suppression of the *orc4-R267A* lethal phenotype by different general transcription factor alleles. Ten-fold serial dilutions of *orc4-R267A* plasmid-shuffle strains without and with additional point mutations in the indicated genes involved in RNA Pol II transcription. Strains were incubated for 4 days at 25 °C. *orc4-* and *toa2-sofa* alleles are included as references. For TBP, we designed *spt15* alleles according to the crystal structure (A86D, R107A) and tested previously-reported alleles that disrupt TFIIA (K133R, Y139A) or TFIIB (E188A) binding [44,45,61,62]. For the rest of the general transcription factors, we tested two *sua7* (TFIIB) alleles (K201I and F289S) [63], a deletion of the non-essential *DST1* gene (TFIIS) [64,65], two *ssl2* (a subunit of TFIIH) alleles (H508R and V552I/E556K) [66,67], and two mutations in *rpo21* (M818I and G1437D) [68].

General transcription factors (GTF), such as TFIIA, promote RNA Pol II transcription initiation through the formation of the preinitiation complex (PIC). There are seven RNA Pol II general transcription factors (GTFs) that participate in this process: TFIIA, TFIIB, TFIID, TFIIE, TFIIF, TFIIH and TFIIS [69]. GTFs promote PIC formation and function by recognizing promoter elements, recruiting RNA Polymerase II, and initiating promoter melting. TFIIS helps RNA Pol II processivity during transcription elongation but can associate with the PIC [70,71].

To test whether any mutation that reduces RNA Pol II transcription suppresses *orc4-R267A* lethality, we introduced a panel of loss-of-function alleles from different GTFs into the *orc4-R267A* plasmid-shuffle strain [30,31,72]. At least one allele from each gene showed significant suppression of the *orc4-R267A* lethal phenotype (Fig 6). The TOA1 allele, *toa1-V251F*, two TBP alleles (*spt15-Y139A* and *spt15-E188A*) and an RNA Pol II allele (*rpo21-G1437D*) were the strongest suppressors. Both TFIIB alleles (*sua7-K201I* and *sua7-F289S)* and a deletion of TFIIS (*DST1::KAN*) showed intermediate suppression. The TFIIH alleles, *ssl2-H508R* and *ssl2-V552I/E556K* were the weakest. Three TBP mutations and one RNA Pol II allele did not show significant suppression. The suppression across all the alleles decreased at 30°C (S6 Fig). These results suggest that suppression reflects a genome-wide reduction in RNA Pol II transcription rather than a Toa2-specific effect. This establishes a connection between transcription and ORC ATPase function.

## Discussion

The Orc4-R267A mutation abolishes ORC’s ATPase activity and is lethal in yeast. However, the protein complex with this mutation retains ORC’s ability to load the Mcm2-7 complex *in vitro*. In this study, we identified six alleles that suppress *orc4-R267A* lethality. Two of the five suppressor mutations in ORC increase the *in vitro* helicase-loading activity of the ATPase-deficient ORC protein while genetic analysis establishes potential connections between the two ATPase sites present in the ORC-Cdc6 complex. Finally, the identification of multiple alleles in RNA Pol II general transcription factors that suppress *orc4-R267A* connects transcription to ORC ATPase function.

We found that the ORC4R protein has a DNA-binding defect in addition to its ATPase defect. In contrast to seven-fold stimulation seen for the wild-type ORC protein, ATP does not stimulate DNA binding of the ORC4R protein (Fig 3B). This defect is consistent with studies of human ORC mutants at the analogous position, which also eliminate ATP-dependent DNA binding [73]. Although this defect could be due to a loss of ATP binding by ORC4R, previous studies found no defect in this function [25]. In addition, ORC4R consistently retains two- to three-fold more helicase-loading activity than the ATP-binding deficient ORCKT (S5C Fig). Thus, although ORC4R has a clear ATP-dependent DNA binding defect, other activities, presumably dependent on ATP hydrolysis, are also impacting its function.

The Orc1 and Orc4 suppressor mutations increase the loading activity of the ATPase-deficient ORC protein (ORC4R) (Fig 3C, S4A and S4B Figs). This finding suggests that these mutations push the level of helicase loading provided by the *orc4-R267A* allele above a level required for viability. Our biochemical studies and available structural models do not reveal an underlying mechanism of this restoration of helicase loading. The effect does not appear to be at the level of intrinsic DNA binding or ATPase activity (Fig 3A and 3B). Given that we measured ORC ATP hydrolysis and DNA binding in the absence of other helicase loading proteins, it is possible that the suppressor mutations alter these functions in a way that only occurs in the context of the full loading reaction. Alternatively, the impact of the suppressors could be downstream and independent of these events (e.g., disassembly of the OCCM).

In contrast to the Orc1 and Orc4 suppressor mutations, the Orc3, Orc5 and Orc6 mutations decrease the loading activity of ORC4R in our experiments. One limitation of our biochemical assays is the absence of chromatinized templates, which is known to impact multiple steps in DNA replication initiation [54]. In addition, ORC participates in the establishment of the nucleosome-free region found at origins of replication [27,74,75]. ORC, via the Orc1 BAH and IDR domains, also recruits chromatin remodelers to promote proper nucleosome array phasing [27] and can evict the H2A/H2B histone dimer in an ATP hydrolysis dependent manner [76,77]. Finally, BAH-independent ORC-nucleosome binding has also been reported [76,78]. Because the Orc3, Orc5, and Orc6 *sofa* mutations did not impact DNA binding or helicase loading, we suggest that they could suppress via chromatin and/or chromatin-remodeler-mediated mechanisms. Although we did not identify chromatin-remodeling genes in our screen, this is consistent with our screen not reaching saturation and the expectation that potentially rare gain-of-function interaction mutations in chromatin remodelers might be required to satisfy the screen.

Our genetic analysis groups the *sofa* alleles into two sets; those that exclusively suppress defects in the Orc1-Orc4 ATPase interface and the *orc1-sofa* allele, which also suppresses alterations in the Cdc6-Orc1 ATPase interface (Fig 1C and 4A). Cdc6 overexpression mirrors the pattern of most *sofa* alleles (suppress *orc4-R267A* and *orc1-K485T*, but not *orc1-R616A*) (Fig 5), suggesting these alleles act similarly, possibly by stabilizing ORC on DNA [4,5,10,79]. The ability of the *orc1-sofa* to also suppress *orc1-R616A,* suggests a different mechanism of suppression for this allele, possibly via events depending on ORC-Cdc6 ATP hydrolysis. Indeed, the ability of *orc1-sofa* to suppress mutations in both the Orc1-Orc4 and Cdc6-Orc1 ATPase sites raises the interesting possibility that these ATPases both stimulate the same event. This is consistent with structural studies showing similar DNA contacts with the Orc1-Orc4 and Cdc6-Orc1 ATPases and suggests inter-ATPase communication [80].

The *toa2-sofa* allele was the only non-ORC allele identified in the screen and, like most *sofa* alleles, suppressed both Orc1-Orc4 ATPase mutations (Fig 1C and 4A). Although not as penetrant as the *sofa* alleles, TBP (SPT15), TFIIB (SUA7), TFIIS (DST1) and RNA Pol II (RPO21) mutations also suppressed the *orc4-R267A* lethal phenotype (Fig 6, S6 Fig) [44,45,59,62–64,66–68,81]. Since multiple GTFs suppressed the lethal phenotype, the *toa2-sofa* activity is likely due to its effects on general RNA Pol II transcription as opposed to a unique interaction with origin licensing factors. Why didn’t we recover more RNA Pol II GTF mutations in our screen? One explanation is that the suppressor activity of these alleles is generally weaker at the temperature our screen was performed at (30 °C) (S6 Fig).

We propose two mechanisms by which the loss-of-function GTF alleles may suppress the *orc4-R267A* lethal phenotype. First, these mutations could reduce collisions between the replisome and transcription machinery during S phase, which can trigger fork collapse, double-stranded breaks, and under replicated DNA [82–84]. Less fork collapse would reduce the need for additional origins of replication to fill in replication gaps that would arise because of these collisions. Second, reduced transcription could lessen transcriptional disruption of helicase loading during G1 [32,36,37,85]. The ORC4R and ORC1K proteins show DNA-binding and helicase loading defects, which would translate to less replication forks *in vivo*. A reduction in genome-wide transcription, including cryptic and pervasive transcription [72,86,87], through GTF loss of function could increase the efficiency of enough origins to overcome the defects of the ORC ATPase alleles. Regardless of the mechanism, our findings reveal a genetic interaction between ORC ATPase function and the general transcription machinery, a relationship that may have broader relevance given that transcription shapes replication initiation zones in mammalian genomes [88–91]. Indeed, because metazoan ORC lacks sequence-specific DNA binding [92,93], the interplay between transcription and origin licensing may play an greater role in determining where replication initiates in these organisms.

## Methods

### Yeast strains and plasmids

All *S. cerevisiae* strains were congenic with W303 (*ade2-1 trp1-1 leu2-3,112 his3-11,15 ura3-1 can1-100*) except the *CDC6* swapper strains (yHR050 and yLM150) which are congenic to S288C. Strain genotypes are summarized in S1 Table. Plasmids are summarized in S2 Table. Oligonucleotides and guide RNAs are summarized in S3 Table.

### Screen for orc4-R267A suppressors

yLM009 was grown in -ADE, -URA medium till 0.5-0.6 OD. Cells were washed with water and resuspended in 100 mM sodium phosphate buffer (pH 7.4). EMS was added to 3% and incubated for 25 min. Cells were washed with 9% (g/mL) sodium sulfate three times and allowed to recover for 3 h in YPD then plated on 5-FOA-containing plates and grown for 5 days. Red single colonies were restreaked on 5-FOA plates. Colonies were restreaked on YPD plates for 2 days and compared to a wild-type reference strain. Strains with similar growth to the starting strain were backcrossed to yLM010 (a *MAT***a** version of the starting strain) and dissected on YPD using an MSM400 (Singer Instruments). Mutant strains that showed 2:2 segregation of the 5-FOA-resistance phenotype were backcrossed a second time. After the second backcross, strains with consistent 2:2 segregation of 5-FOA-resistance phenotype were selected for pooled-linkage analysis and whole-genome sequencing [40].

### Pooled-linkage analysis and whole-genome sequencing

Pooled-linkage analysis was carried out as described [40,94]. After the second tetrad dissection of each mutant strain, ten spore colonies with the suppressor phenotype were pooled and genomic DNA was prepared [95]. This process was repeated with 10 spore colonies from the same dissection that lacked the phenotype. Illumina libraries were generated from 1 ng of genomic DNA using Nextera XT (Illumina) and sequenced on an Illumina HiSeq2000 using a single-end 40 nt read.

Fastq files were aligned to the *S. cerevisiae* S288C genome assembly R64 using bwa (0.7.17) [96]. Sequences were sorted and indexed using Samtools [97]. Potential causative mutations were identified using a custom-made python script and the following criteria: 1) More than 10 reads mapped to the base; 2) more than 90% of reads have the same nucleotide and 3) the nucleotide was different between the wild-type and suppressor samples.

### Generation of point mutations

Yeast strains were transformed using the standard LiOAc/PEG transformation protocol [98]. Point mutations were introduced into the endogenous locus using Cas9-mediated gene editing based on previously published protocols [42,43]. Colonies were genotyped using PCR [99]. Guide RNA sequences targeted in transformations are listed in S3 Table.

### Plasmid-shuffle assay

Yeast strains were grown overnight under non-selective conditions for the URA3-containing plasmid (YEP + 2% Glu or 2% Galactose). Ten-fold serial dilutions were plated and incubated for 2-3 days on the indicated media at the indicated temperatures.

### Purification of ORC, Mcm2-7, and Cdc6

ORC, Mcm2-7 and Cdc6 were purified as described previously [16,18,100]. In strains co-expressing wildtype V5-tagged Orc4 and untagged Orc4-R267A (yLM001-006), Flag-purified ORC contains a mixture of wildtype and mutant complexes. To separate them, we incubated the peak Flag-elution fractions with anti-V5 agarose beads (Sigma-Aldrich) overnight at 4°C, depleting wildtype ORC and enriching for mutant complexes. After this incubation, the beads were pelleted by centrifugation and separated from the supernatant. Downstream steps were performed as described previously.

### Electrophoretic-mobility shift assays

The radioactive *ARS1* DNA template was prepared by PAGE-purified restriction-enzyme digested products of plasmid p*ARS1*/WT. The ends were filled with dNTPs and radioactive dATP (5 µCi) and this reaction was purified using Amersham MicroSpin G-50 Columns (27533001). The assays were performed as described previously [51] except that 2.5% (v/v) glycerol was included in the PAGE gel. For ATP-free reactions, 0.25 U of apyrase (NEB: M0398S) was added before the addition of the ORC protein. All DNA templates are WT *ARS1* unless stated otherwise. All reactions used ATP unless indicated otherwise. All images were collected on a GE Amersham Typhoon Biomolecular Imager.

### ATPase assay

ATP hydrolysis assays were performed as previously described [26]. A 90 min-timepoint sample without ORC was used to subtract background signal from all samples. All images were collected on a GE Amersham Typhoon Biomolecular Imager.

### Helicase-loading Assay

Reconstituted helicase-loading assays were performed as described previously [16]. All reactions contained 3mM ATP unless otherwise indicated. For experiments in which one reaction contained less than 3 mM ATP, an ATP-regenerating system consisting of 10 µg creatine kinase (Sigma: 10127566001) and 50 mM phosphocreatine (Sigma: P7936) was included in all reactions. All DNA templates are WT *ARS1* unless indicated otherwise. All images were collected on a GE Amersham Imager 680.

### Image quantification and data visualization

ImageJ (v1.54g) was used to quantify images from electrophoretic-mobility shift, ATPase, and helicase-loading assays. R 4.4.3 was used to align and plot the multiple sequence alignments in S2 and S3 Figs. R 4.4.3 was used to generate the plots in Figs 3A, S4A, and S4B.

## Acknowledgements

We thank Marko Lõoke for his help with the variant-calling analysis of the suppressor screen. During manuscript preparation, Claude (Anthropic) was used to suggest revisions to sentence wording and to assist with code development. All AI-assisted outputs were reviewed, edited, and validated by the authors, who take full responsibility for the accuracy, originality, and integrity of the manuscript, analyses, and code.

## Supporting information

**S1 Fig. Hydroxyurea (HU) sensitivity and temperature sensitivity of the *sofa*-allele with different ORC ATPase mutations.**

Ten-fold serial dilutions of (A and B) *ORC4, orc4-R267A*, (C and D) *ORC1, orc1-K485T and orc1-R616A* plasmid-shuffle strains with additional *sofa* allele. Strains were incubated 2-3 days at 30°C in 50mM HU or 2-3 days at 37°C.

**S2 Fig. Conservation of Orc1, Orc3 and Orc4 *sofa* alleles residues.**

Multiple sequence alignments were generated using MUSCLE algorithm and similarity was calculated according to the BLOSUM62 scoring matrix. Residues are colored according to their chemical properties (e.g., hydrophobic, polar, positively/negatively charged). Black boxes enclose the alignment column for the mutated residue in the *sofa* alleles. A) Orc1. B) Orc3. C) Orc4.

**S3 Fig. Conservation of Orc5, Orc6 and Toa2 *sofa* alleles residues.**

Multiple sequence alignments were generated using MUSCLE algorithm and similarity was calculated according to the BLOSUM62 scoring matrix. Residues are colored according to their chemical properties (e.g., hydrophobic, polar, positively/negatively charged). Black boxes enclose the alignment column for the mutated residue in the *sofa* alleles. A) Orc5. B) Orc6. C) Toa2.

**S4 Fig. Additional biochemical characterization of ORC4R*^sofa^* proteins.**

A) Representative *in vitro* helicase-loading assay for WT, ORC4R, and ORC4R^Orc4-sofa^ proteins across a potassium glutamate titration (250, 300, and 350 mM), with quantification below (n = 3). B) Representative *in vitro* helicase-loading assay for WT, ORC4R, and ORC4R^sofa^ proteins at 350 mM potassium glutamate (compared to 300 mM in Fig 3C), with quantification below (n = 3).

**S5 Fig. Additional biochemical characterization of the ORC4R protein.**

A) Comparison of DNA binding of WT ORC, and ORC4R with and without the Orc4-*sofa* mutation, and ORC1K in electrophoretic-mobility shift assay (EMSA). Binding in the presence of ATP or ATPγS is shown. B) Sequence specificity of ORC and ORC4R DNA binding. EMSA was used to measure wild-type ORC and ORC4R DNA binding to WT *ARS1* or *ARS1* lacking the strongest ORC binding site (ACS-). C) Titration of WT ORC, ORC4R, and ORC1K proteins in a helicase-loading assay (0.3 mM ATP). D) Salt titration and sequence-specific loading of the WT and ORC4R protein (R267A). Sequence specificity is tested using a WT *ARS1* and an *ARS1* lacking the two strongest ORC binding sites (ACS-/B2-).

**S6 Fig. Suppression of the *orc4-R267A* lethal phenotype by general transcription factor alleles at 30.**

Ten-fold serial dilutions of *ORC4* or *orc4-R267A* plasmid-shuffle strains with additional point mutation in the indicated gene involved in RNA Pol II transcription. Strains were incubated for 4 days at 30. Strains are the same as Fig 6.

**S1 Table. List of *Saccharomyces cerevisiae* strains used in this study**

**S2 Table. List of plasmids used in this study**

**S3 Table. List of oligonucleotides and guide RNAs used in this study**

